# Stromal and Neuronal Sources of Slit2/3 Ligands in the Adult Pancreas Exhibit Distinct Expression Patterns Independent of Robo2 Receptor Expression in the Islet

**DOI:** 10.64898/2026.05.15.725534

**Authors:** Matthew R. Wagner, Nicolas G. Pintozzi, Bjorn M. Schoff, Marissa I. Gold, Rachel H. Kasper, Nina G. Steele, Barak Blum

**Affiliations:** Department of Cell and Regenerative Biology, University of Wisconsin-Madison, School of Medicine and Public Health, Madison, WI, United States; Division of gastroenterology and hepatology, Department of Internal Medicine, University of Cincinnati, College of Medicine, Cincinnati, OH, United States

## Abstract

Pancreatic islets regulate blood glucose homeostasis. Although islet architecture is stable under homeostatic conditions, increased metabolic demand drives compensatory islet expansion. In mice, islets are organized as a β cell core surrounded by a mantle of α and δ cells. The formation of islet architecture during development requires expression of Roundabout receptors 1 and 2 (Robo1/2) in endocrine cells and of Slits 2 and 3 (Slit2/3) from islet-extrinsic sources. Furthermore, expression of Robo2 in endocrine cells is required to maintain islet architecture in the adult mouse. However, the cellular sources of Slit2/3 in the adult pancreas and their expression dynamics during islet expansion remain unknown. Here, we identify distinct stromal populations, including fibroblasts and pericytes, as well as neurons within intrapancreatic ganglia, as the sources of Slit2/3. We further show that Slit3 expression is increased in Ob/Ob mice, and that SLIT2 expression is elevated in stromal cell populations of humans with type 2 diabetes. The expression of neither Slit2 nor Slit3 was affected by deletion of Robo2 in β cells. Together, these findings define the cellular origins of Slit2/3 and their expression dynamics in the adult pancreas, supporting a potential role for Slit signaling in the diabetic islet microenvironment.

## INTRODUCTION

The pancreatic islets of Langerhans function to maintain euglycemia by secreting hormones from endocrine cells. Murine islets are comprised of three main endocrine cell types (α, β, δ, secreting glucagon, insulin, and somatostatin, respectively) and are arranged with a mantle of α and δ cells surrounding a core-like structure of β cells^1^. Human islets are comprised of the same aforementioned cell types, but are arranged more sporadically^1^. Despite these architectural differences, the β cells in both humans and mice preferentially organize to form homotypic β cell-to-β cell connections^2^. These homotypic β cell-to-β cell interactions are important for the coordinated and synchronous release of islet hormones in response to stimuli, thereby regulating systemic blood glucose levels^2–4^. Defects in murine islet architecture, as a consequence of genetic or pharmacological experiments, have been associated with aberrant regulation of blood glucose^5,6^. Additionally, human patients with type 2 diabetes (T2D) histologically present with disrupted islet architecture^1,7^.

Slit-Robo signaling is involved in regulating islet architecture and islet function^8–12^. Deletion of both *Robo1* and *Robo2* (*Robo1/2*) in embryonic β cells during islet development, or of *Robo2* alone in adult β cells, is sufficient to interrupt the formation and maintenance of mature islet architecture, respectively^8,10^. Whole-body deletion of *Slit2* and *Slit3* (*Slit2/3*) together, but not deletion of *Slit1*, disrupts islet morphogenesis during mouse development, and function of either Slit2 or Slit3 alone rescues the phenotype, thus showing that Slit2 and Slit3 play redundant roles in the formation of islet architecture *in vivo*^9^. Finally, the expression of *Robo* receptors in the islet is diminished during islet compensatory expansion in obesity and diabetes^8^, suggesting dynamic involvement of the Slit-Robo pathway in islet morphological changes in the adult.

*Slit2* and *Slit3* ligands have been shown to be expressed by the pancreatic mesenchyme components throughout pancreas organogenesis and in later adult stages^9,11,13,14^. However, the specific cell types within the mesenchyme population that express Slit2/3 in the adult pancreas, and the extent to which this expression is dynamic during islet compensatory expansion are unknown. Here, we set out to spatially characterize the expression of *Slit2/3* in the adult mouse and human pancreas and their expression dynamics during islet compensatory expansion.

## RESULTS

### *Slit2* and *Slit3* are expressed by pancreatic stromal and neuronal cell types

To determine the cell types that express *Slit2/3* in mature pancreata, we combined RNA fluorescence *in situ* hybridization (RNAscope) for *Slit2/3* with immunofluorescence using specific markers for a broad variety of pancreatic cell types (Figure 1A-I). We thus looked for *Slit2/3* expression in acinar cells (α amylase), endothelial cells (CD31), leukocytes (CD45), pericytes (NG2), fibroblasts (Vimentin), glial cells (S100β), and neurons (β-Tubulin III, NeuN, and ChAT). Across all cell types, we identified two predominant expression patterns of *Slit2/3*: one that is low expression of either *Slit2* or *Slit3,* and one that is high expression of both *Slit2* and *Slit3*. The low expression of either *Slit2* or *Slit3* alone was limited to the leukocytes, pericytes, fibroblasts, and glial cells, with less than ten percent of each cell type expressing either gene (Figure 1J-K). The high expression of both *Slit2* and *Slit3* together was reliably confined to neurons, with one hundred percent of neurons expressing each gene (Figure 1L). These low and high expression patterns in mature pancreata are consistent with results obtained by another group^14^.

**Figure 1.**
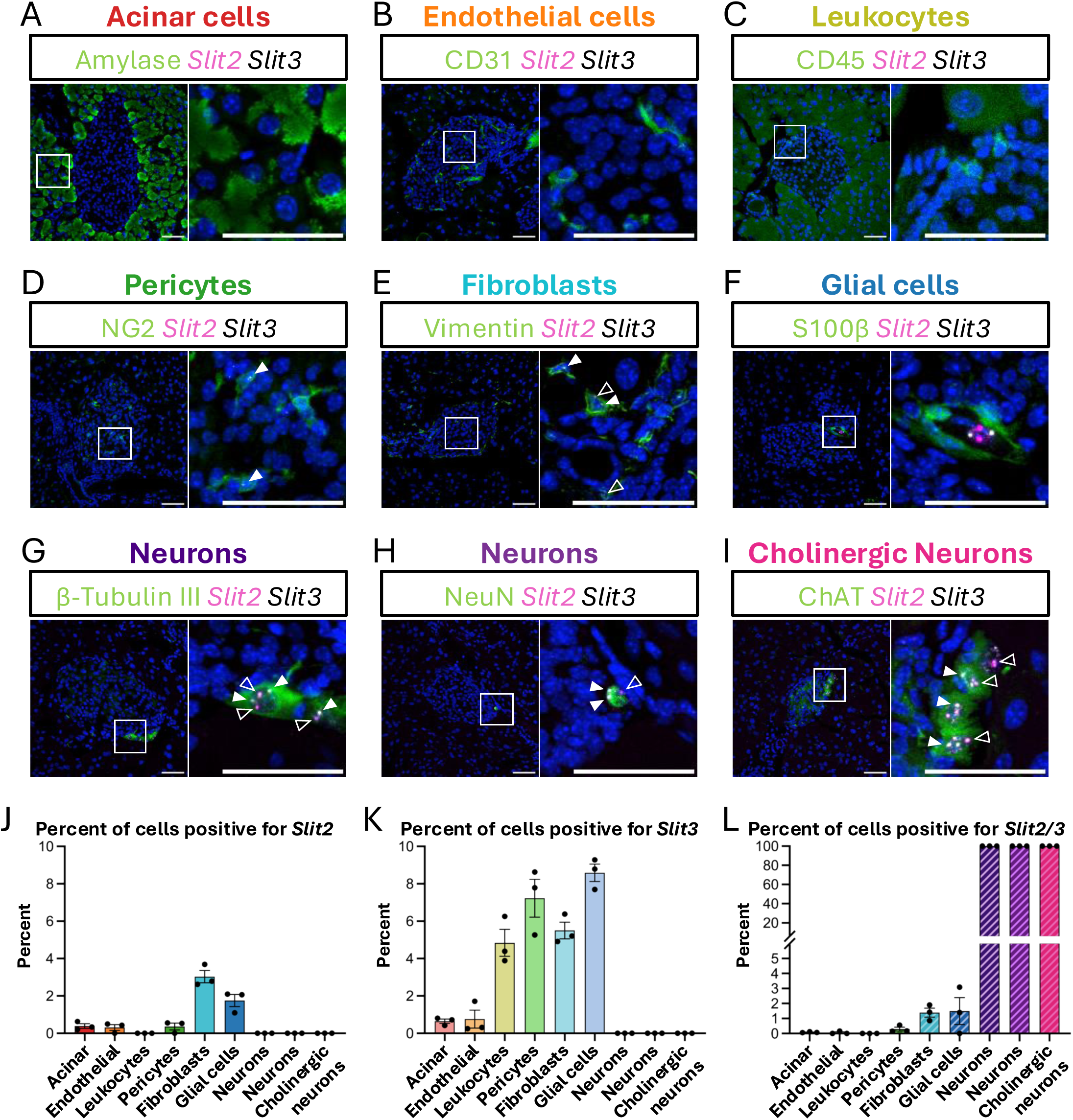
*Slit2* and *Slit3* are expressed by stromal and neuronal cell types. **(A-I)** mRNA expression of *Slit2* (magenta) and *Slit3* (white) in **(A)** acinar cells, **(B)** endothelial cells, **(C)** leukocytes, **(D)** pericytes, **(E)** fibroblasts, **(F)** glial cells, and **(G-I)** neurons. **(J-L)** Percentages of cell types that express **(J)** *Slit2*, **(K)** *Slit3*, and **(L)** *Slit2/3*. Nuclei are stained with DAPI. Scale bars correspond to 50μm. N=3 mice, 16-29 images per mouse.

### Spatial distribution of *Slit2/3*-expressing cells surrounding islets

To test the spatial distribution of *Slit2/3*-expressing cells surrounding pancreatic islets, we analyzed the *Slit2/3* RNAscope images by segmenting them to generate spatially distinct, 20μm-wide concentric bands surrounding islets (Figure 2A). We detected 359,563 cells across all images and regions, of which 7,044 were *Slit+*, equating to roughly 2% of all detected cells. Within the *Slit+* cells, we identified the proportion of cells that express *Slit2* only, *Slit3* only, and both *Slit2* and *Slit3* together, with the most abundant being *Slit3+* at 62.76%, followed by *Slit2+* at 26.78% and *Slit2+/3+* at 10.45% (Figure 2B). In all combinations of *Slit2/3* expression, we found that there is a greater percentage of *Slit+* cells proximal to islets, in the 0-20μm band, when compared with the more distal, 20-40μm, 40-60μm, 60-80μm, and 80-100μm bands (Figure 2C). To identify the contributions of *Slit* expression in each band, we looked at the proportion of cells expressing *Slit2* and *Slit3* and found that the percentage of *Slit3*-expressing cells is significantly higher than *Slit2/3*-expressing cells in the Extra-islet 1 and Extra-islet 3 bands (Figure 2D). We further looked at the mRNA expression level (puncta/cell) of *Slit2* and *Slit3* in each band and found that there is a similar gradient, with cells proximal to islets having slightly higher expression of *Slit2* and *Slit3* (Figure 2E).

**Figure 2.**
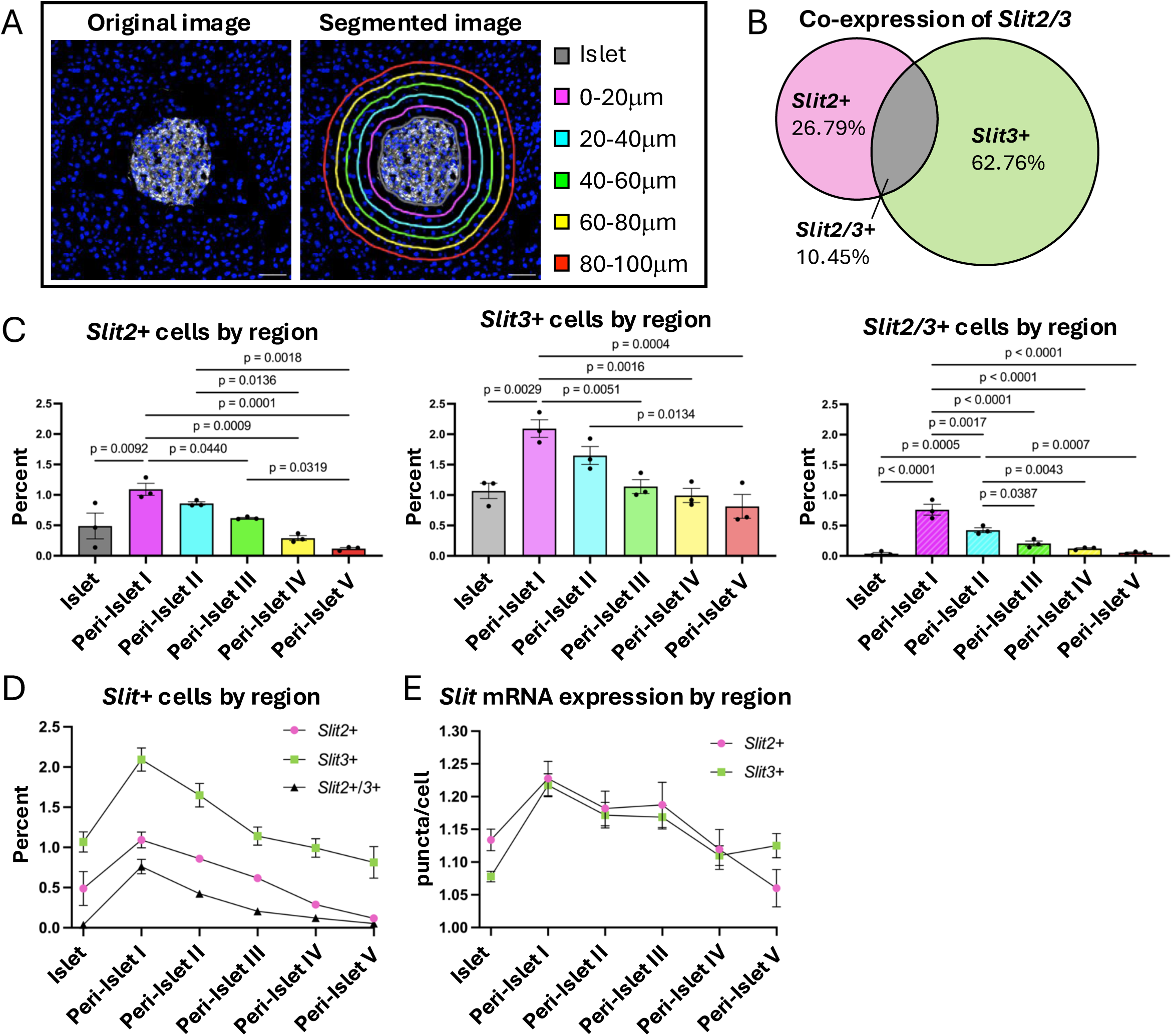
Spatial distribution of Slit2/3-expressing cells surrounding islets. **(A)** A schematic of segmented regions. **(B)** Percentages of all Slit-expressing cells that express *Slit2*, *Slit3*, or *Slit2/3*. **(C-D)** Percentages of cells that express *Slit2*, *Slit3*, and *Slit2/3* within each region. **(E)** Average expression of *Slit2* and *Slit3* in Slit-expressing cells within each region. N=3 mice, with 144-208 images per mouse, analyzed using one-way ANOVA for multiple comparisons.

### Expression of *Slit3*, but not *Slit2*, is increased in pancreatic stromal cells of *Ob/Ob* mice

The expression of *Robo2*, a canonical receptor for the Slit ligands, is decreased in pancreatic islets of obese and diabetic humans and mice^8,15,16^. To a different extent, the expression of *Slit2* mRNA has been shown to change in acute pancreatitis and pancreatic ductal adenocarcinoma^14^. To test whether *Slit2/3* expression is dynamic under diabetogenic stress, we performed RNAscope using pancreatic tissue from lean wild-type and obese leptin-null and diabetic (*Ob/Ob*) mice, and Vimentin as a cell marker, since fibroblasts were seen to express both *Slit2* and *Slit3* in previous results (Figure 3A). Upon quantification, the percentage of Vimentin+ cells that expressed *Slit2* or *Slit3* remained unchanged in the *Ob/Ob* mice; however, the expression level of *Slit3* was significantly increased in the Vimentin+ cells of *Ob/Ob* mice (Figure 3B). We further analyzed the non-immunolabeled, unclassified cells in the same tissues and observed a significant increase in the percentage of cells that express *Slit3*, as well as the level of *Slit3* expression in the *Ob/Ob* mice (Figure 3C-D). No changes in the percentage of cells expressing *Slit2* or the expression level of *Slit2* were found between *Ob/Ob* and control mice.

**Figure 3.**
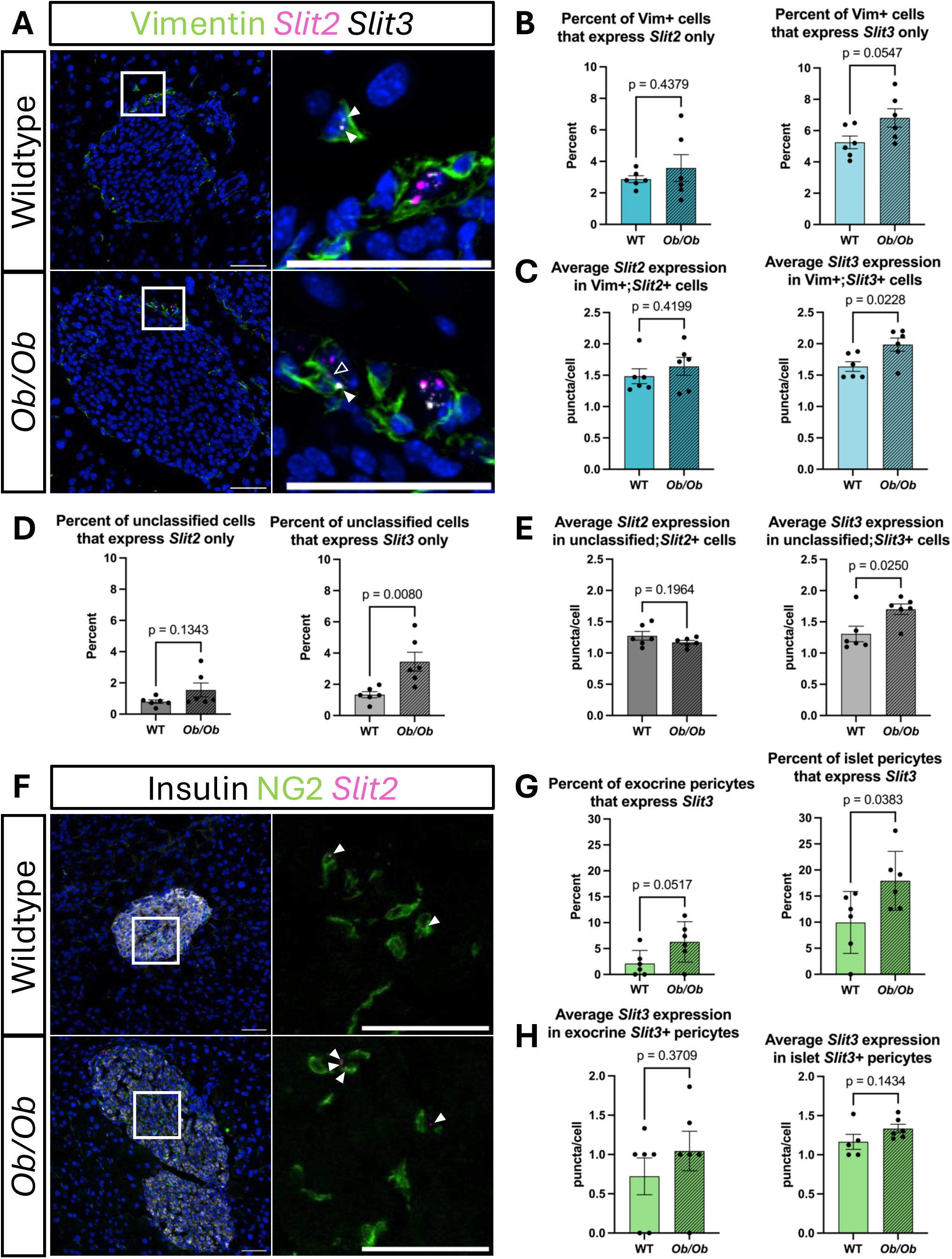
*Expression of Slit3, but not Slit2, is increased in pancreatic stromal cells of Ob/Ob mice*. **(A)** mRNA expression of *Slit2* (magenta) and *Slit3* (white) in fibroblasts (green). **(B)** Percentages of fibroblasts that express *Slit2* and *Slit3* in wild-type and *Ob/Ob* mice. **(C)** Average *Slit2* and *Slit3* expression in fibroblasts in wild-type and *Ob/Ob* mice. **(D)** Percentages of unclassified cells that express *Slit2* and *Slit3* in wild-type and *Ob/Ob* mice. **(E)** Average *Slit2* and *Slit3* expression in unclassified cells in wild-type and *Ob/Ob* mice. **(F)** mRNA expression of *Slit3* (magenta) in pericytes (green) of wild-type and *Ob/Ob* mice. **(G)** Percentages of exocrine and islet pericytes that express *Slit3*. **(H)** Average *Slit3* expression in exocrine and islet pericytes in wild-type and *Ob/Ob* mice. Nuclei are stained with DAPI. Scale bars correspond to 50μm. N=6 mice per group, with 15 to 28 images per mouse, analyzed via unpaired t-test.

We hypothesized that pericytes, one of the predominant *Slit3-*expressing cell types from our previous results, would change *Slit3* expression levels in obesity and diabetes as well. To test the hypothesis that *Slit3* expression would change in pericytes, we performed RNAscope with NG2 as a cell marker of pericytes and insulin to identify islets (Figure 3F). To determine whether pericytes located within the islets or in the surrounding exocrine tissue express *Slit3* at differing levels, we spatially segmented the islets from the surrounding 75μm of exocrine tissue, to capture intra-islet and extra-islet pericytes, respectively. We found that the percentage of pericytes expressing *Slit3* is greater inside islets compared to pericytes found in the outside the islet, regardless of genotype, with 3-4 times more pericytes expressing *Slit3* in islets than pericytes from the surrounding tissue (Figure 3G). Between genotypes, the percentage of pericytes that express *Slit3* is greater in *Ob/Ob* mice than compared to control animals (Figure 3G). Quantification of *Slit3* mRNA expression in pericytes did not reveal significant differences between *Ob/Ob* and control mice in either intra-islet or extra-islet pericytes (Figure 3H).

### Expression of Slit2/3 does not change in the intrapancreatic ganglia of *Ob/Ob* mice

To assess *Slit2/3* expression in the other major cell type that expresses *Slit2/3*, the neurons, we analyzed RNAscope images containing neurons of both wildtype and *Ob/Ob* mice (Figure 4A). Upon quantification, we did not observe changes in the percentages of cells that express *Slit2*, *Slit3*, or *Slit2* and *Slit3* (Figure 4B). Additionally, neither the expression of *Slit2* nor *Slit3* were changed in the neurons (Figure 4C).

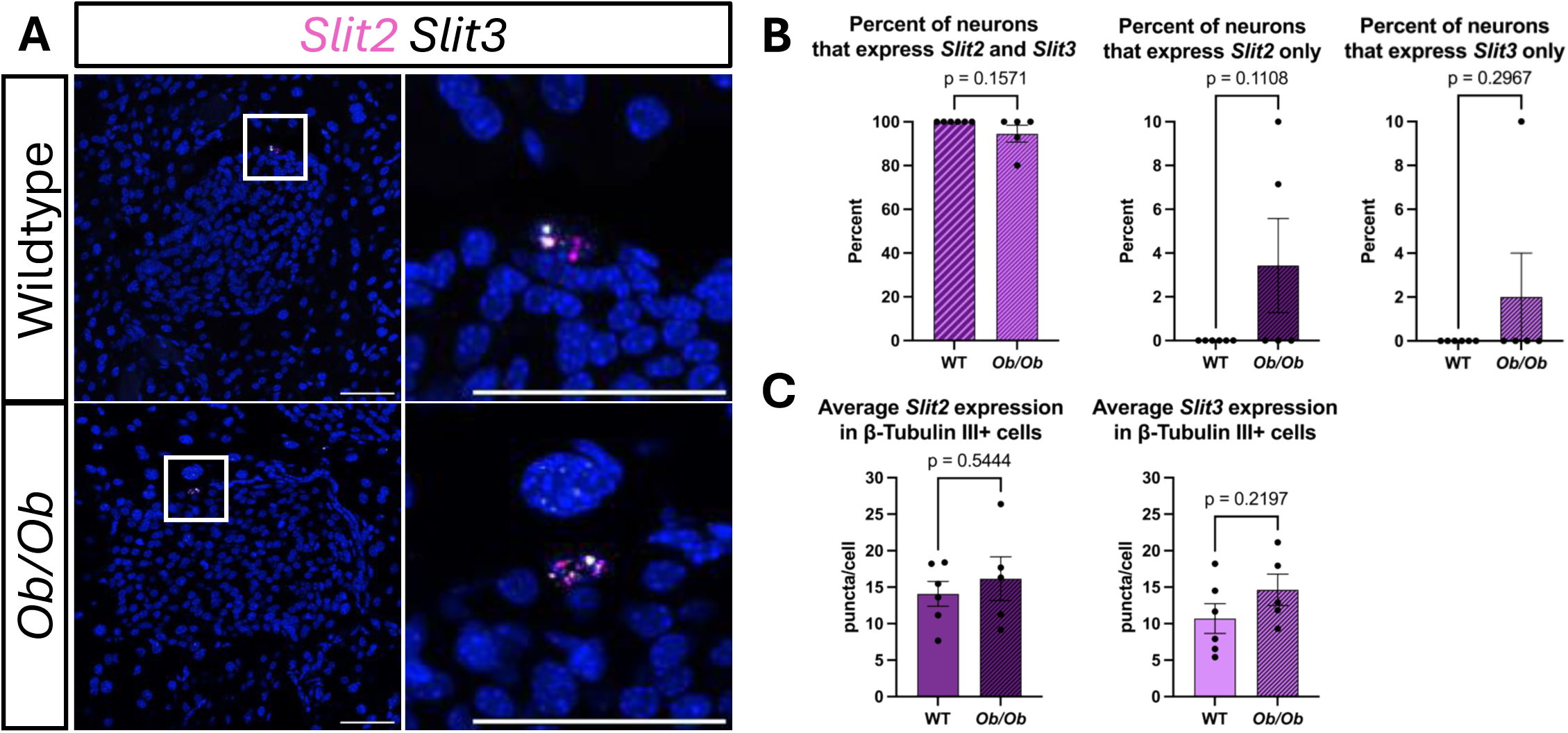

### Expression of extra-islet *Slit2/3* is not dependent on the expression of *Robo2* in β cells

To investigate whether *Slit* expression is dependent on the reduction of *Robo2* expression seen in obesity and diabetes^8^, we generated β cell-specific *Robo2*-knockout mice (Figure 5A). We performed RNAscope for *Slit2/3* using pancreatic tissue from wild-type (Robo2^+/+^) and β cell-specific *Robo2*-knockout (Robo2^fl/fl^) mice, and Insulin as a cell marker to identify the islets (Figure 5B). Upon quantification of *Slit2/3* expression, we did not observe any changes in the percent of cells that express *Slit2* or *Slit3* (Figure 5C), nor did we observe changes in the expression of *Slit2/3* mRNA puncta per cell across genotype (Figure 5D). Thus, the removal of *Robo2* expression in β cells does not induce expression of *Slit2/3* in the surrounding cells.

**Figure 5.**
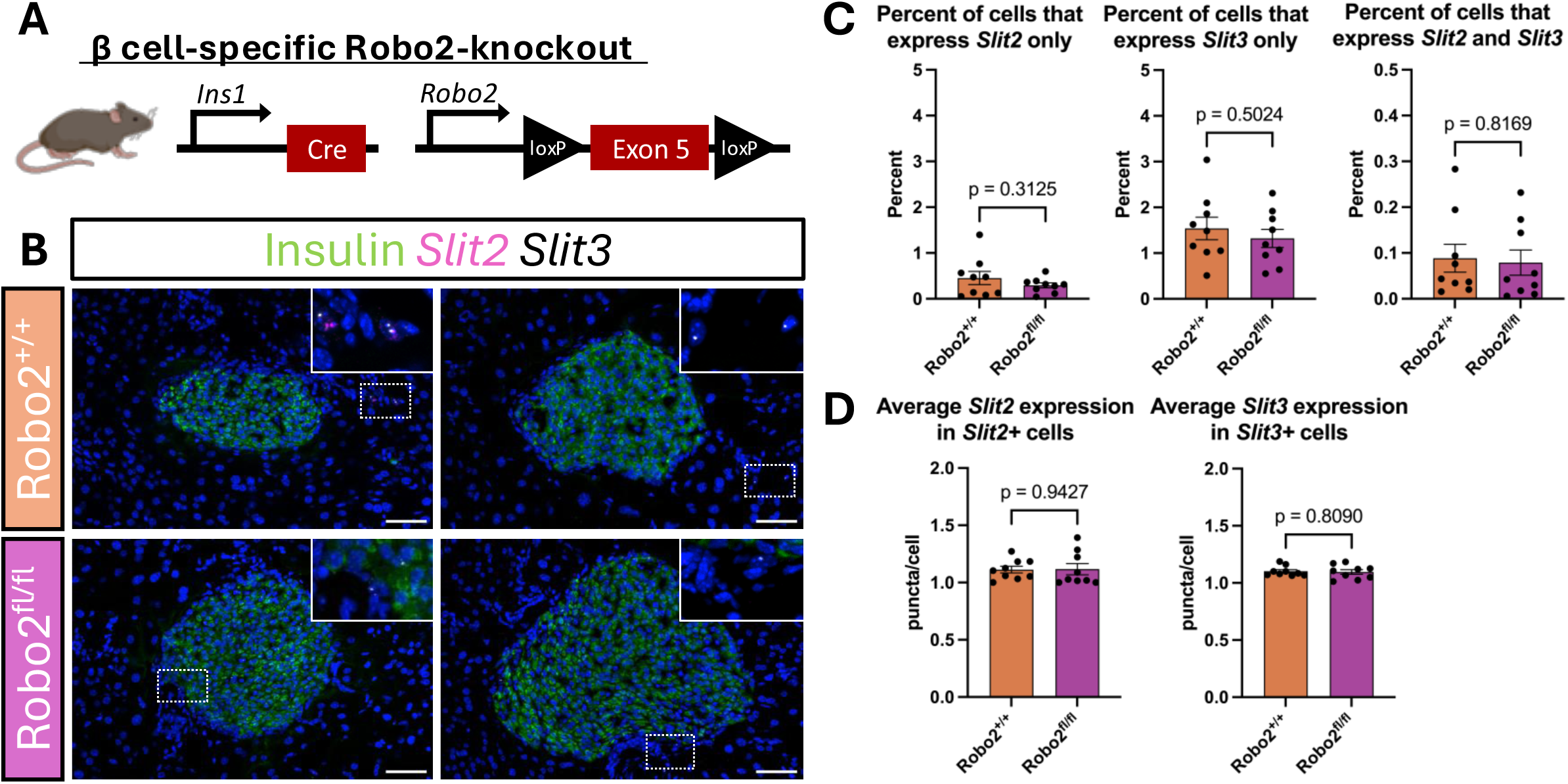
Expression of extra-islet *Slit2/3* is not dependent on the expression of *Robo2* in β cells. **(A)** Schematic of alleles used for generating β-cell-specific Robo2-knockout mice. **(B)** mRNA expression of *Slit2* (magenta) and *Slit3* (white) in and around islets (Insulin, green) of control and experimental mice. **(C)** Percentages of cells that express *Slit2* and *Slit3* in control and experimental mice. **(D)** Average *Slit2* and *Slit3* expression in cells of control and experimental mice. Nuclei are stained with DAPI. Scale bars correspond to 50μm. N=9 mice per group, with 7 to 31 images per mouse, analyzed via unpaired t-test.

### Human *SLIT2* and *SLIT3* are expressed by stromal cell populations and their expression levels are altered in T2D

To assess the expression of *Slit2/3* in the human pancreas, we examined a previously published single-cell RNAseq dataset that utilizes samples from non-diabetic (n=29) and T2D (n=17) human donors^17^. We identified 14 cell types across all samples, with *SLIT2* and *SLIT3* expression primarily localized to the mesenchyme cell clusters (Figure 6A-B, and Supplementary Figure 1A). *SLIT2* expression is predominantly within the mesenchyme 1 population, which is defined by its expression of *COL6A3* (Supplementary Figure 1A). *SLIT3* is largely expressed by the mesenchyme 2 population, which is defined by its expression of *RGS5* (Supplementary Figure 1A), and further has moderate expression in the mesenchyme 1 population as well. To test whether the expression of human *SLIT2/3* changes with diabetogenic conditions like that of mouse *Slit2/3*, we examined at the expression of *SLIT2* and *SLIT3* in the Mesenchyme 1 (Mes1) and Mesenchyme 2 (Mes2) clusters of non-diabetic and T2D samples. Converse to the mouse data, we did not observe changes in *SLIT3* expression in neither the Mes1 nor the Mes2 cluster. We did, however, observe increased *SLIT2* expression in T2D samples within the Mes1 cluster, where *SLIT2* is primarily expressed (Figure 6C). In both mesenchyme 1 and 2 clusters, 22.53% of cells are *SLIT+*, with the most abundant being *SLIT3+* at 51.62%, followed by *SLIT2+* at 43.32% and *SLIT2/3+* at 5.06% (Figure 6D). To dissect the specific populations of cells within the two mesenchyme clusters, we subset and re-clustered the mesenchymal cells and identified 8 cell types (Figure 6E, Supplementary Figure 1B). Similar to the mouse data, we found that *SLIT2* is mainly expressed by the fibroblast populations and that *SLIT3* is more highly expressed in pericytes but is also expressed to a lower degree in the fibroblast populations (Figure 6F).

**Figure 6.**
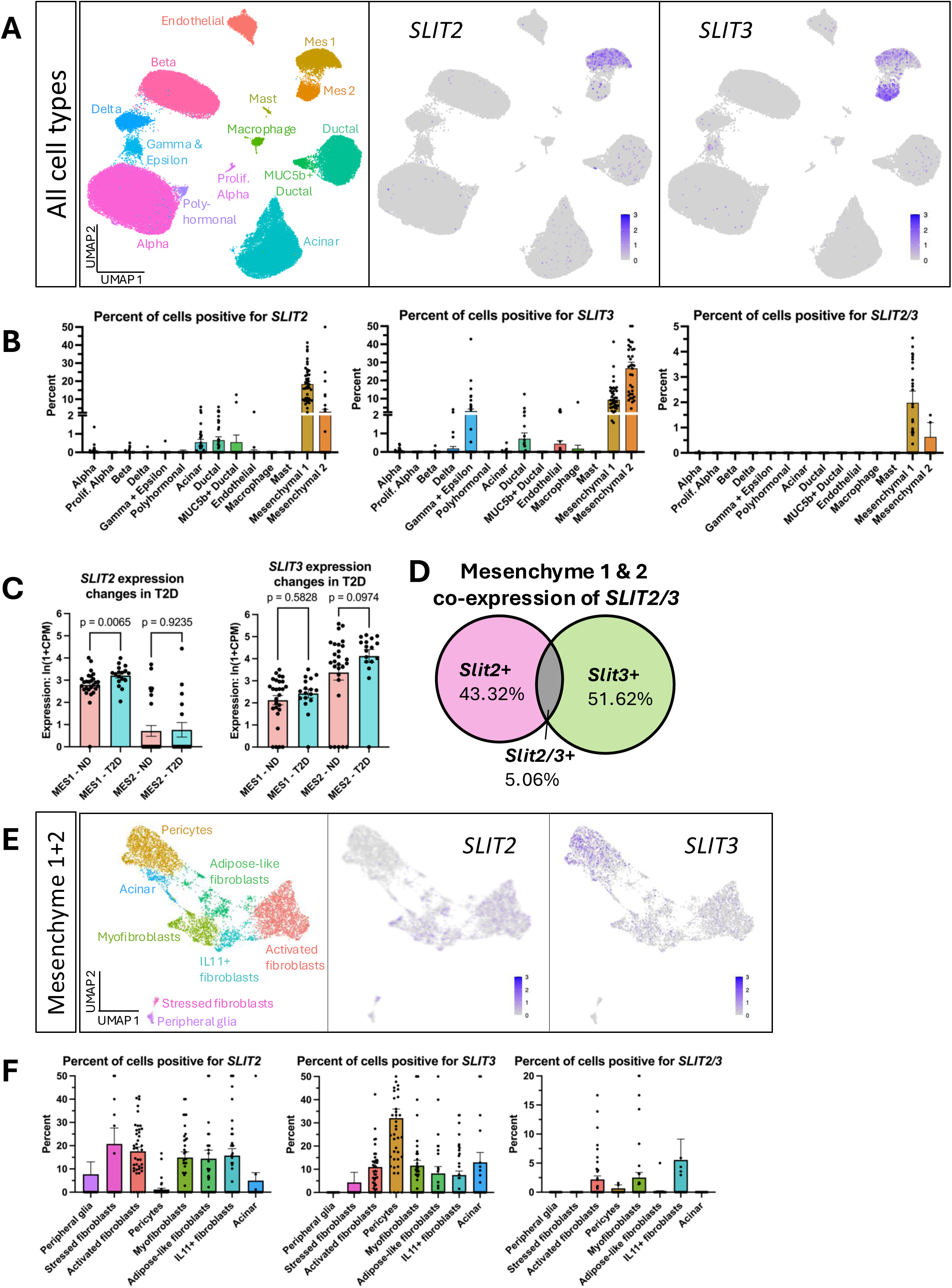
Single-cell RNA-seq analysis of non-diabetic and T2D human pancreas samples. **(A)** UMAP plot displaying 14 cell clusters identified via gene marker expression, and expression of *SLIT2* and *SLIT3*. **(B)** Percentages of cell types that express *SLIT2*, *SLIT3*, and *SLIT2/3*. **(C)** Expression of *SLIT2* and *SLIT3* in the mesenchymal clusters between non-diabetic and T2D samples. **(D)** Percentages of all Slit-expressing cells that express *Slit2*, *Slit3*, or *Slit2/3*. **(E)** Subclustered UMAP plot displaying 8 cell types within the Mesenchyme 1 and Mesenchyme 2 cell populations and expression of *SLIT2* and *SLIT3*. **(F)** Percentages of cell types that express *SLIT2*, *SLIT3*, and *SLIT2/3*. N=29 for ND and N=17 for T2D, analyzed via Mann-Whitney U test.

## DISCUSSION

In this study, we set out to spatially characterize the expression of *Slit2/3* in the adult mouse and human pancreas and their expression dynamics during islet compensatory expansion. We identify stromal and neuronal cells as the primary sources of Slit2/3 within the adult pancreas, supporting a model in which Slit2/3 ligands are supplied by the islet microenvironment rather than endocrine cells themselves^11,14,18^. Notably, *Slit2/3* expression is heterogeneous among the stromal cell populations, whereas the parasympathetic neurons strongly and uniformly express both *Slit2* and *Slit3*, confirming that cellular niches differently contribute to local Slit availability^11^. These findings are consistent with prior reports of mesenchymal Slit expression during pancreas development and histological observations of low *Slit2* expression within pancreatic mesenchymal cells and high *Slit2* expression in other cells, likely neurons, found adjacent to pancreatic islets^14,18^. Together, these data support that stromal and neural cells are the sources of Slit signaling to the Robo-expressing pancreatic islet cells across developmental and adult contexts.

Extending these observations to human tissue, we find that fibroblasts and pericytes represent the principal sources of *SLIT2* and *SLIT3*, respectively, reinforcing the concept that non-endocrine cell types predominantly contribute to Slit ligand production. Although neuronal populations were not captured in the single-cell dataset, the similarity between murine and human stromal expression demonstrates that the microenvironment origin of Slit signaling is broadly conserved. This cross-species agreement strengthens the interpretation that the function of Slit ligands as paracrine cues within the islet niche may be conserved.

Our data further demonstrate that Slit expression is dynamically regulated in obesity and diabetes. Consistent with prior observations of increased *Slit2* expression in acute pancreatitis and pancreatic ductal adenocarcinoma^14^, we observe elevated Slit ligand expression in diabetic conditions, including increased *Slit3* in the *Ob/Ob* murine model and *SLIT2* in human datasets. This divergence in Slit expression between humans and mice may reflect species-specific regulation of the genes in response to metabolic challenges or differences in extracellular matrix organization and stromal signaling cues as a result of the different basement membrane organization between the two species^19^. More broadly, these findings suggest that Slit signaling is responsive to changes in metabolic demand and tissue architecture.

Importantly, genetic ablation of *Robo2* did not alter *Slit2/3* expression, indicating that ligand production may be independent from the expression of *Robo2*. The independence from receptor expression supports a model in which Slit ligands are regulated extrinsically, through microenvironment or systemic factors, and may act as adaptive signals during islet remodeling to support endocrine cell function^11,12^.

Together, these findings establish Slit2/3 as dynamically regulated, microenvironment-derived signals within the pancreas and raise several important directions for future study. A key next step will be to define the functional consequences of *in vivo* Slit signaling in adult islets, particularly whether Robo activation regulates β cell survival, maturation, or transcriptional identity through downstream gene networks. Additionally, our identification of both neuronal and mesenchymal sources of Slit ligands suggests that these cells may provide distinct and potentially non-redundant inputs to islets. Dissecting the relative contributions of these sources will provide important insight into how islet architecture and function are maintained or disrupted in diabetes through microenvironment changes, as neuronal-derived Slit signaling may allow for more rapid responses to changes in systemic or autonomic input, whereas the mesenchymal-derived Slit signals may reflect a sustained response to systemic changes and lead to remodeling of the extracellular matrix or islet architecture. Given the observed changes in Slit and Robo expression across metabolic states, such studies may reveal a role for the Slit-Robo pathway in coordinating adaptive versus maladaptive responses in diabetes progression.

## METHODS

### Animals and Tissue Preparation

Animal experiments were conducted in accordance with the University of Wisconsin-Madison Institutional Animal Care and Use Committee (IACUC) under protocol number M005221. C57BL6/J (Strain #000664) and C57BL6-Lep^Ob^/J (*Ob/Ob*; Strain #:000632) mice were obtained from Jackson Laboratory and were aged 8-10 weeks old. All *Ob/Ob* mice had an ambient blood glucose >300mg/dL before tissue collection. The Robo2 βKO model contained the Ins1-Cre, Robo2-flx, and Rosa26-LSL-H2B-mCherry alleles, which have been previously described (Refs.^20–22^). Robo2 control mice were Ins1-Cre^Tg/0^; Robo2^+/+^; Rosa26-LSL-H2B-mCherry^Tg/0^ and Robo2 βKO mice were Ins1-Cre^Tg/0^; Robo2^flx/flx^; Rosa26-LSL-H2B-mCherry^Tg/0^. Adult mice, at least 12 weeks of age, were maintained on normal chow diet (Envigo, Teklad 2018). Pancreata of mice were collected and prepared as fixed-frozen slides as previously described^10^.

### RNAscope and Confocal Imaging

RNAscope experiments were performed using the Multiplex Fluorescent V2 Assay Kit (Cat. No. 323100; Advanced Cell Diagnostics (ACD)) and the RNA-Protein Co-Detection Ancillary Kit (ACD, Cat. No. 323180), following the manufacturer’s protocols (ACD, 323100-USM and MK 51–150 TN). Murine probes for *Slit1* (#502491), *Slit2* (#449691), *Slit3* (#542771-C2) from ACD were used. TSA Plus Cyanine 3.5 (Cat. No. NEL763001KT), Cyanine 5 (Cat. No. NEL745001KT), and Cyanine 5.5 (Cat. No. NEL766001KT) fluorophores from Quanterix (formerly Akoya Biosciences) were diluted 1:500 in TSA Buffer (ACD, Cat. No. 322809) and used to target the RNA probes. Primary antibodies used for immunostaining were guinea pig anti-Insulin, 1:4 (Agilent, IR002); mouse anti-Glucagon, 1:200 (Sigma-Aldrich, G2654); rabbit anti-Somatostatin, 1:100 (Phoenix Pharmaceuticals, G-060-03); rabbit anti-Amylase 1:50 (Cell Signaling Technology, 3796); rat anti-CD31, 1:50 (BD Pharmingen, 550274); rat anti-CD45, 1:10 (BD Pharmingen, 550539); rabbit anti-NG2, 1:200 (Sigma-Aldrich, AB5320); rabbit anti-Vimentin, 1:200 (Abcam, AB92547); rabbit anti-S100β, 1:50 (Invitrogen, PA5-78161); rabbit anti-βTubulin III, 1:1,000 (BioLegend, 802001); rabbit anti-NeuN, 1:100 (Cell Signaling Technology, 24307); rabbit anti-ChAT, 1:25 (Invitrogen, PA5-29653). Secondary antibodies used for immunostaining were AlexaFluor 488 (anti-rabbit, anti-rat, and anti-guinea pig) from Jackson ImmunoResearch, and were diluted 1:500 in the ACD Co-Detection Blocking Solution along with DAPI at 1:10,000 (Sigma, 9542) to counterstain nuclei. Slides were imaged using a 40X oil objective lens on a Nikon AXR confocal microscope housed in the University of Wisconsin Optical Imaging Core (Grant #1S10O34394-01).

### QuPath Analysis, Image Segmentation in Fiji, and Statistics

Image analysis was performed using maximum-intensity-projection images of z-series acquired on the UWOIC Nikon AXR. Briefly, nuclei were detected using Watershed cell detection, and composite classifiers were generated to simultaneously identify cell types and mRNA puncta based on pixel intensity. Annotated images were assessed and manually corrected in cases of erroneous pixel-intensity detections, such that as high background signals of the immunostaining. Detection measurements of mRNA count per cell were exported and analyzed using RStudio to generate average puncta per cell and percent of cells expressing the mRNA. The data are presented as average ± standard error of the mean. All statistics were performed with Prism GraphPad 10 unless otherwise stated. P-values <0.05 were considered significant.

### Single-cell RNAseq Analysis

We utilized scRNA-seq data of human isolated pancreatic islets made available by the Human Pancreas Analysis Program (RRID:SCR_016202) and integrated by members of the Gaulton lab. Briefly, the Gaulton lab used Cell Ranger 6.0.1 (10× Genomics) to align HPAP data to the human genome and removed cells with less than 500 expressed genes per cell and less than 15% mitochondrial reads. Ambient RNA was removed using SoupX 1.6.1^23^, and doublets were identified and removed by Scrublet 0.2.3^24^. Upon downloading the integrated Seurat object, additional filtering was performed to exclude cells with more than 5,000 expressed genes, more than 30,000 RNA counts, or less than 500 RNA counts. All type 1 diabetes and type 1 diabetes autoantibody-positive cells were removed from downstream analysis. The remaining non-diabetic and T2D cells were log-normalized using a scale factor of 10,000, and the top 2000 most variable genes were identified using the variance-stabilizing-transformation method. Data were scaled and principal component analysis was completed within Seurat 5.2.1^25^. Batch integration was performed using Harmony 1.2.4^26^, incorporating library, tissue source, and chemistry as covariates. The harmony-corrected embeddings and 16 principal components were used for UMAP dimensionality reduction plots and clustering. For gene-level visualization, raw counts were aggregated at the donor level within each cell type to generate pseudobulk profiles prior to CPM normalization and ln(1+CPM) transformation.

## Acknowledgements

We thank members of the Blum laboratory, particularly Cyrus R. Sethna, for valuable discussion and comments on the manuscript. We are also grateful to Lance Rodenkirch and the UW-Madison Optical Imaging Core for help with imaging. This work was funded by R01DK121706 from the NIDDK to BB.

## Author Contributions

Conceptualization: M.R.W. and B.B.; Methodology: M.R.W.; N.G.S., and B.B.; Investigation and Analysis: M.R.W., N.G.P., B.M.S., M.I.G., and R.H.K.; Writing original draft: M.R.W. and B.B.; Reviewing and Editing: All Authors; Funding acquisition: B.B.; Supervision: B.B.

---

**Figure S1.**
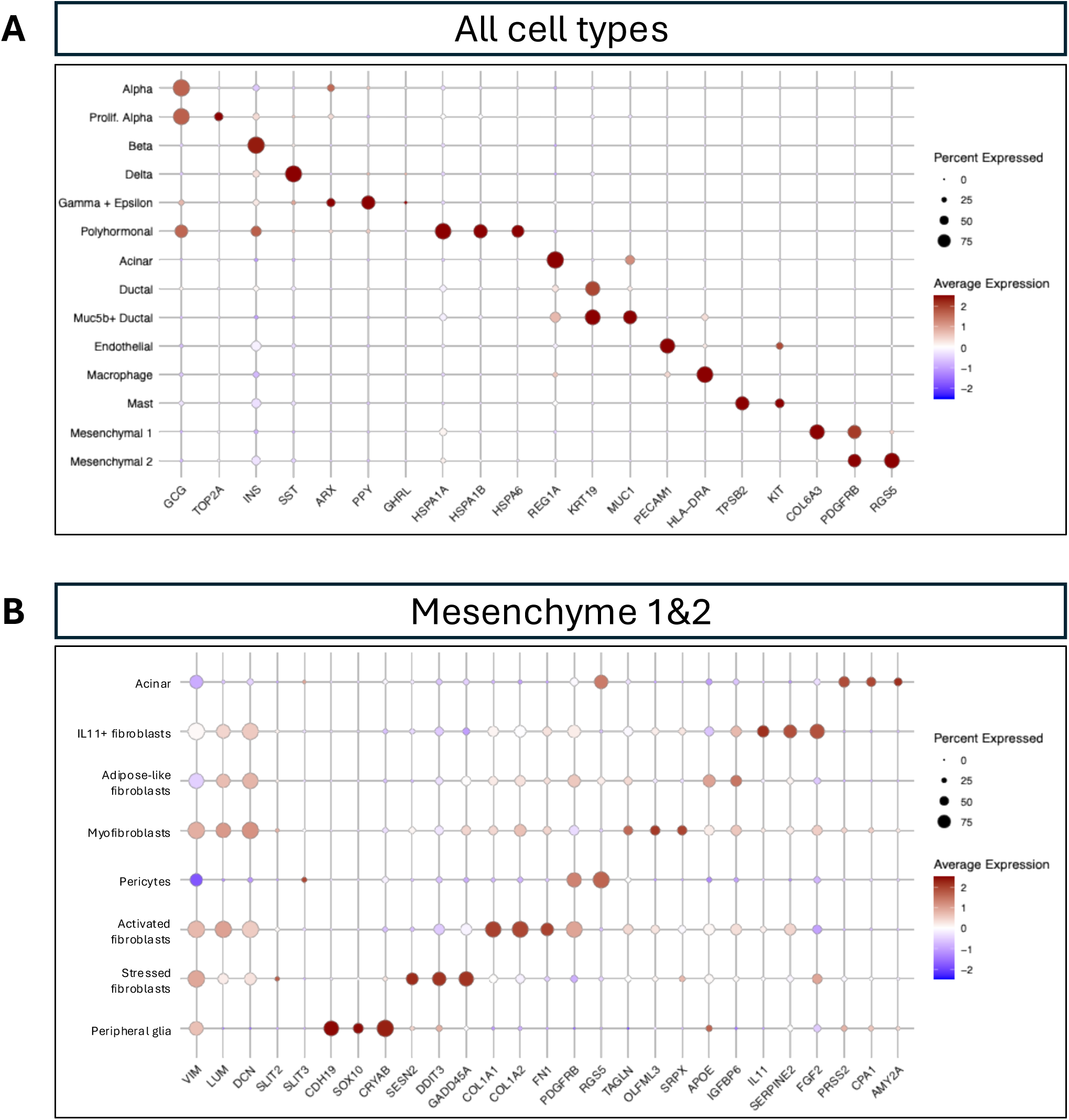
Single-cell RNA-seq cluster identification. **(A)** Bubble plot displaying 14 cell clusters identified via gene marker expression. **(B)** Subclustered bubble plot displaying 8 cell types within the Mesenchyme 1 and Mesenchyme 2 cell populations.

## REFERENCES

1. Steiner, D.J., Kim, A., Miller, K. & Hara, M. Pancreatic islet plasticity: interspecies comparison of islet architecture and composition. Islets 2, 135–145 (2010).

2. Hoang, D.T., et al. A conserved rule for pancreatic islet organization. PloS one 9, e110384 (2014).

3. Adams, M.T., et al. Reduced synchroneity of intra-islet Ca(2+) oscillations in vivo in Robo-deficient beta cells. eLife 10(2021).

4. Hoang, D.T., Hara, M. & Jo, J. Design Principles of Pancreatic Islets: Glucose-Dependent Coordination of Hormone Pulses. PloS one 11, e0152446 (2016).

5. Adams, M.T., Waters, B.J., Nimkulrat, S.D. & Blum, B. Disrupted glucose homeostasis and glucagon and insulin secretion defects in Robo betaKO mice. FASEB journal: official publication of the Federation of American Societies for Experimental Biology 37, e23106 (2023).

6. Borden, P., Houtz, J., Leach, S.D. & Kuruvilla, R. Sympathetic innervation during development is necessary for pancreatic islet architecture and functional maturation. Cell reports 4, 287–301 (2013).

7. Brereton, M.F., Vergari, E., Zhang, Q. & Clark, A. Alpha-, Delta- and PP-cells: Are They the Architectural Cornerstones of Islet Structure and Co-ordination? The journal of histochemistry and cytochemistry: official journal of the Histochemistry Society 63, 575–591 (2015).

8. Adams, M.T., Gilbert, J.M., Hinojosa Paiz, J., Bowman, F.M. & Blum, B. Endocrine cell type sorting and mature architecture in the islets of Langerhans require expression of Roundabout receptors in beta cells. Scientific reports 8, 10876 (2018).

9. Gilbert, J.M., Adams, M.T., Sharon, N., Jayaraaman, H. & Blum, B. Morphogenesis of the Islets of Langerhans Is Guided by Extraendocrine Slit2 and Slit3 Signals. Mol Cell Biol 41, e0045120 (2021).

10. Waters, B.J., Birman, Z.R., Wagner, M.R., Lemanski, J. & Blum, B. Islet architecture in adult mice is actively maintained by Robo2 expression in beta cells. Developmental biology 505, 122–129 (2024).

11. Cozzitorto, C., et al. A Specialized Niche in the Pancreatic Microenvironment Promotes Endocrine Differentiation. Developmental cell 55, 150–162 e156 (2020).

12. Yang, Y.H., Manning Fox, J.E., Zhang, K.L., MacDonald, P.E. & Johnson, J.D. Intraislet SLIT-ROBO signaling is required for beta-cell survival and potentiates insulin secretion. Proceedings of the National Academy of Sciences of the United States of America 110, 16480–16485 (2013).

13. Escot, S., Willnow, D., Naumann, H., Di Francescantonio, S. & Spagnoli, F.M. Robo signalling controls pancreatic progenitor identity by regulating Tead transcription factors. Nature communications 9, 5082 (2018).

14. Pinho, A.V., et al. ROBO2 is a stroma suppressor gene in the pancreas and acts via TGF-beta signalling. Nature communications 9, 5083 (2018).

15. Xin, Y., et al. RNA Sequencing of Single Human Islet Cells Reveals Type 2 Diabetes Genes. Cell metabolism 24, 608–615 (2016).

16. Fadista, J., et al. Global genomic and transcriptomic analysis of human pancreatic islets reveals novel genes influencing glucose metabolism. Proceedings of the National Academy of Sciences of the United States of America 111, 13924–13929 (2014).

17. Elgamal, R.M., et al. An Integrated Map of Cell Type-Specific Gene Expression in Pancreatic Islets. Diabetes 72, 1719–1728 (2023).

18. Torres-Cano, A., et al. Spatially organized cellular communities shape functional tissue architecture in the pancreas. Sci Adv 11, eadx5791 (2025).

19. Otonkoski, T., Banerjee, M., Korsgren, O., Thornell, L.E. & Virtanen, I. Unique basement membrane structure of human pancreatic islets: implications for beta-cell growth and differentiation. Diabetes, obesity & metabolism 10 Suppl 4, 119–127 (2008).

20. Thorens, B., et al. Ins1(Cre) knock-in mice for beta cell-specific gene recombination. Diabetologia 58, 558–565 (2015).

21. Lu, W., et al. Disruption of ROBO2 is associated with urinary tract anomalies and confers risk of vesicoureteral reflux. Am J Hum Genet 80, 616–632 (2007).

22. Blum, B., et al. Reversal of beta cell de-differentiation by a small molecule inhibitor of the TGFbeta pathway. eLife 3, e02809 (2014).

23. Young, M.D. & Behjati, S. SoupX removes ambient RNA contamination from droplet-based single-cell RNA sequencing data. Gigascience 9(2020).

24. Wolock, S.L., Lopez, R. & Klein, A.M. Scrublet: Computational Identification of Cell Doublets in Single-Cell Transcriptomic Data. Cell systems 8, 281–291 e289 (2019).

25. Hao, Y., et al. Dictionary learning for integrative, multimodal and scalable single-cell analysis. Nature biotechnology 42, 293–304 (2024).

26. Korsunsky, I., et al. Fast, sensitive and accurate integration of single-cell data with Harmony. Nat Methods 16, 1289–1296 (2019).

